# Excitability regulation in the dorsomedial prefrontal cortex during sustained instructed fear responses: a TMS-EEG study

**DOI:** 10.1101/277806

**Authors:** Gabriel Gonzalez-Escamilla, Venkata C. Chirumamilla, Benjamin Meyer, Tamara Bonertz, Sarah von Grothus, Johannes Vogt, Albrecht Stroh, Oliver Tüscher, Raffael Kalisch, Muthuraman Muthuraman, Sergiu Groppa

## Abstract

**Background:** Threat detection is essential for protecting individuals from precarious situations. Early studies suggested a network of amygdala, limbic regions and dorsomedial prefrontal cortex (dmPFC) involved in fear processing. Excitability regulation in the dmPFC might be crucial for physiological fear processing, while an abnormal excitability pattern could lead to mental illness. Non-invasive paradigms to measure excitability regulation during fear processing in humans are missing.

**Methods:** We adapted an experimental approach of excitability characterization using electroencephalography (EEG) recordings and transcranial magnetic stimulation (TMS) over the dmPFC during an instructed fear paradigm to dynamically dissect its role in fear processing. Event-related (ERP) and TMS-evoked potentials (TEP) were analyzed to trace dmPFC excitability in healthy young volunteers (n = 40, age = 27.6 ± 5.7 years, 22 females). Moreover, we linked the excitability regulation patterns to individual structural MRI-derived properties of gray matter microstructural integrity of the fear network.

**Results:** An increased cortical excitability was demonstrated in the threat (T) condition in comparison to no-threat (NT) as showed by increased amplitude of evoked potentials. Furthermore, TMS over the dmPFC induced markedly increased evoked responses during T condition in relation to NT. Moreover, we found that the structural integrity of the dmPFC and the amygdala predict excitability regulation patterns as measured by ERP and TEP during fear processing.

**Conclusions:** We describe the dynamic range of excitability regulation in dmPFC during fear processing. The applied paradigm can be used to non-invasively track response abnormalities to threat stimuli in healthy subjects or patients with mental disorders.

## Introduction

The dorsomedial prefrontal cortex (dmPFC) is involved in working memory, attention, emotion regulation and further distinct mental functions. Its role in threat processing has been repeatedly postulated [1, 2]. Sustained fear situations are bounded to a well-defined stressor that will occur with some predictability in a short time window [3]. In instructed fear paradigms, a state of fear can be elicited by a cue when there is a contingency between it and a potentially dangerous stimulus. Previous functional (f)MRI studies have shown that evaluation of fearful stimuli lead to an activation of the dmPFC [2]. If excitatory or inhibitory mechanisms are involved or how a regulation of cortical excitability in the dmPFC during fear processing occurs is still unknown. These phenomena, however, play a crucial role for adaptive behavior in threat situations and their dysfunction could lead to the development of neuropsychiatric disorders.

Event-related potential (ERP) analysis is an effective method to address neural processing and cortical excitability. Time-locked synchronous neural activity can be depicted as well during fear processing. One of the characterized ERP components to threat exposures is P100, which reflects the non-conscious processing of presented cues. Conscious processing is linked to sustained attention and increasing use of processing resources [4] and involves medial prefrontal cortex [5]. Fear responsiveness can be indexed by presence of the P300 and the longer-lasting late positive potential (LPP) components. LPP has been as well uniquely linked to memory encoding and storage during fear processing [6]. The LPP amplitude increases to threat stimuli; functional imaging studies showed activation of further nodes of the “fear network” (i.e. amygdala) interrelated to LPP magnitude. Electrophysiological LPP measures are therefore considered a viable marker of fear processing [7, 8] and likely reflect summated neuronal activity of different regions conforming the fear network, given its sensitivity to a variety of task instructions and its long duration which appears not to habituate over repeated presentations of stimuli [9]. Specific causal manipulation trough optogenetics in animal studies or transcranial magnetic stimulation (TMS) in humans could facilitate delimitation of the specific role of network nodes. Of note, the increased positivity of the LPP starting ∼300-500 ms after stimuli presentation extends well beyond 1000 ms [10] and its duration is stimulus-guided, reflecting a continued physiological process related to attentional fear processing [11, 12]. However, despite the importance of LPP to fear processing, its neural substrate is still not clear.

Here, we address dmPFC excitability by analyzing the LPP component through simultaneous TMS and EEG recordings during fear processing. Therefore, we adapted an instructed fear paradigm, where a conditioned stimulus (CS), called the CS+, predicts an unconditional fearful stimulus (US), while the other (the CS-) its absence. This paradigm has been previously applied to map the neural networks engaged in instructed fear showing an involvement of amygdala, insular cortex, the dmPFC, and the anterior cingulate cortex (ACC) [2, 13, 14]. During instructed fear the amygdala and the dmPFC are considered regulatory nodes for network responses to threat [2, 15, 16]. The dmPFC has been suggested to convey excitability regulation states during sustained fear events, via synchronized activity with other brain structures of the fear network [17]. Recent technical advances allow the non-invasive assessment of synchronized brain activity by TMS-EEG recordings at the brain surface level [18-21] and provide the opportunity to track physiological aspects of fear processing at the cortical level while also measuring long-range synchronization and excitability properties within the involved network nodes. In order to dissect and improve the EEG-based moderate spatial resolution for the characterization of the involved network, we link TMS-EEG responses to structural properties of the studied network as derived from MRI. Thereby, a precise temporal and spatial characterization of the dmPFC-guided fear processing is achieved.

## Methods

### Participants

In total, forty healthy young subjects were enrolled for the study. Twenty subjects participated in the designed main experiment (11 female; mean age ± SD: 26.8 ± 4.7 years). A second group of twenty healthy young subjects was used for a control experiment (11 female, mean age ± SD = 28.3 ± 6.6 years). All participants signed an informed consent, approved by the local ethical committee at the University Medical Center Mainz.

### MRI data acquisition

Prior to the instructed fear paradigm each participant underwent an MRI scan protocol. All MRI scans were acquired in a Siemens Trio 3 Tesla scanner (Siemens Medical Solutions, Erlangen, Germany). The T1 structural protocol was a magnetization-prepared rapid gradient-echo (MPRAGE) with the following parameters: repetition time [TR] = 1900 ms; echo time [TE] = 2.54 ms; inversion time [IT] = 900ms; pixel bandwidth = 180; acquisition matrix = 320×320; flip angle = 9°; pixel spacing = 0.8125 x 0.8125 mm; slice thickness = 0.8 mm.

### Instructed fear paradigm

Before starting the experiment, the experimenter explicitly instructed all participants about the fear paradigm. During the sessions, an adapted version of the [Raczka, Mechias [22]] instructed fear paradigm was applied, as previously described [23]. In the threat condition (T), a circle (conditioned stimulus, CS+) was presented, followed in 33% of the cases by a painful electrodermal stimulation (unconditioned stimulus, US) to the back of the right hand (Fig. 1); alternatively, in the no threat condition (NT), a square (unconditioned stimulus, CS-) was presented without any threat stimuli association. The two visual cues (CS+ and CS-) were presented in a pseudorandomized order for 5 seconds, separated by a 5-10 second presentation of a black fixation cross on a white background (ITI).

**Figure 1.**
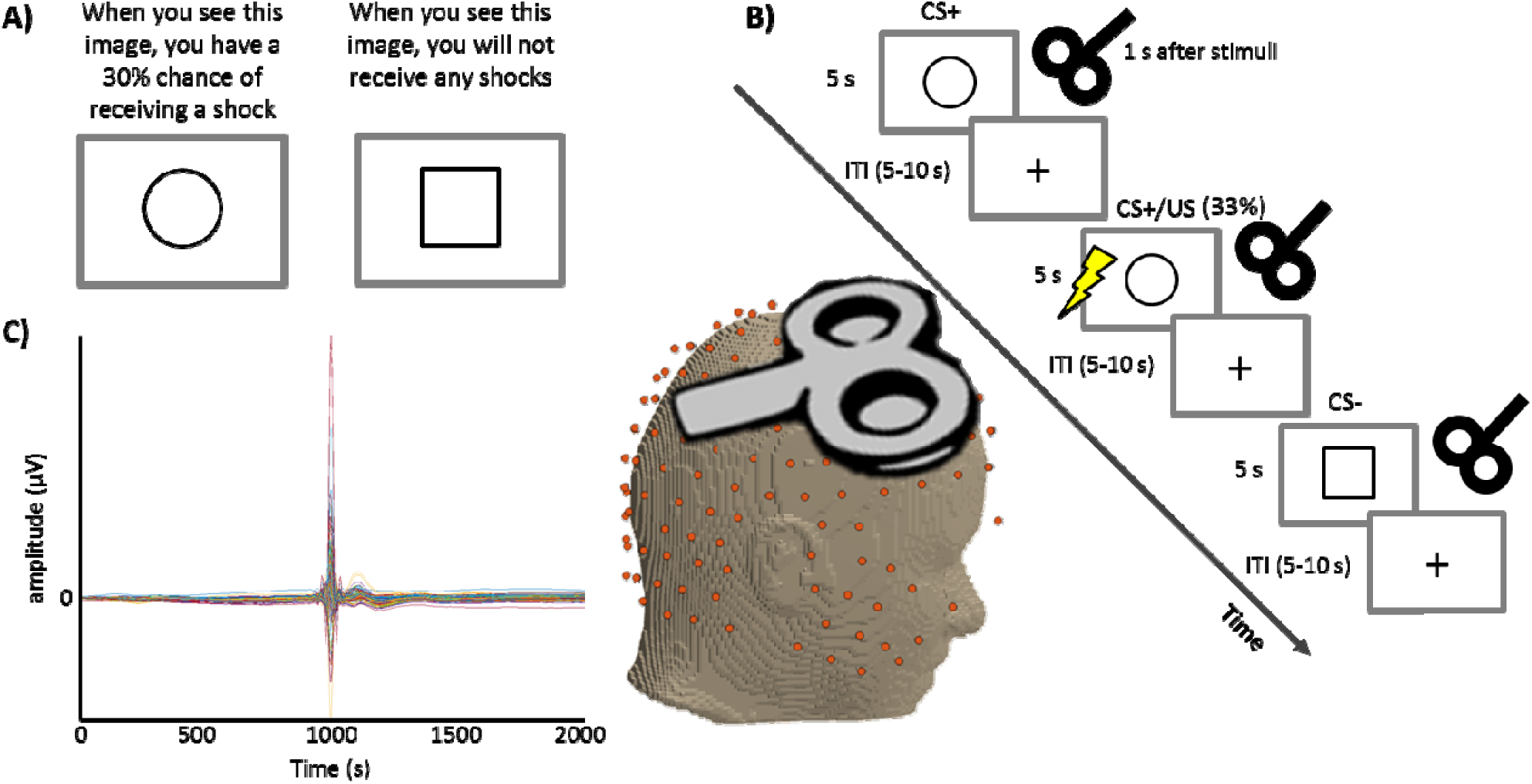
Experimental design. (A) Prior to the experimental phase, participants were informed about paradigm contingencies. (B) Schematic of the fear conditioning task. Three continuous blocks of trials were performed. In each block, one figure (the conditioned stimulus, or CS+) was paired with a shock (the unconditioned stimulus, or US) 33% of the time, whereas a second figure (the CS-) was never paired with a shock. Images were presented for 5 s, followed by a 5-10-second inter-stimulus interval (ITI). The figure represents the pseudorandom trial orders used during the experiment with the TMS stimuli applied 1 s after cue presentation in the case of the TMS experiments.(C) Butterfly plot from one subject showing an example of the acquired EEG data with the TMS pulse visible.

The painful electrical stimuli (US) consisted of square wave pulses of 2 ms each, generated by a DS7A electrical stimulator (Digitimer) and were delivered through a surface electrode situated on the back of the right hand. Prior to the experiment, participant-specific painful stimulus intensity was determined by rating increasing stimulus intensities on a scale form 0 (no pain) to 10 (very painful). An intensity corresponding to pain 7 was used during the experiment.

At the end of every paradigm session the participants reported the amount of acquired fear, referring to their last encounter with each of the two visual cues (scaled to %, 0%= no anxiety, 100% = very anxious). These fear rating scales were accompanied by the caption: “How much fear did you experience while looking at this figure?” There were no time constraints for providing ratings.

### EEG recordings and TMS experiment

EEG signals were recorded continuously during experiments using a high-density (256-channel) EEG recording system (EGI Netstation, Eugene, sampling rate: 250 Hz, impedances: ≤50 kOhms). To establish the intensity of the TMS stimulation (Magstim 200, Magstim Co., Whitland, Dyfed, UK), we first determined the resting motor threshold, defined as the minimum stimulus intensity at which the TMS pulse induced at least five motor evoked potentials (MEP) in ten consecutive trials [19]. MEPs were recorded on the left hand (contralateral hand to the TMS stimulation) abductor pollicis brevis muscle, using a tendon-belly arrangement. During the instructed fear paradigm, we applied the TMS pulses with intensity of 110% of the resting motor threshold [19]. The right dmPFC was target as defined in the individual MR images using a neuronavigation system (Localite, Sank Augustin) to the MNI coordinates ([10 12 58]) delimited in a previous fMRI activation study (Meyer et al., unpublished). The MNI coordinates were registered and transformed to the subject-specific MRI using the SPM8 software (http://www.fil.ion.ucl.ac.uk/spm). Based on previous findings [4, 10, 23], TMS was applied 1000 ms after each visual cue presentation (TMS experiment). A total of 36 TMS pulses were analysed for each condition. Further, as control for the TMS experiment, we applied the same instructed fear paradigm and recorded EEG signals without the addition of TMS. This experiment is referred as no-TMS experiment.

### EEG signal processing and analysis

First, 25 ms of TMS-related artefact (5 ms before and 20 ms after the TMS pulse) was removed from the EEG data. Signals were then processed to account for line-noise and linear trends, channels with high amplitudes over long time periods found during visual inspection were deleted on trial-by-trial and scalp topology maps were used to further identify any remaining channels with artefacts to reject them, noisy channels were finally interpolated. All EEG analyses on the remaining channels were performed using a combination of FieldTrip (http://www.fieldtriptoolbox.org/) and in-house already published scripts [24].The artefact-free EEG data from both experiments was low-pass filtered with a cut-off frequency of 35 Hz and baseline corrected using the 500 ms prior to the visual cue. Event-related (ERP) and TMS-evoked potentials (TEPs) were then computed to identify peaks of activity at the corresponding time intervals of 0 to 1000 ms for ERPs and 1000 to 2000 ms for TEPs. The activity was considered a peak when at least 3 continuous points (12 ms) of the ERP waveform (on both sides) had smaller values. The amplitude at every peak was computed. Furthermore, the difference in amplitudes between threat (CS+ trials without the actual electric shock, US) and no-threat conditions (T-NT) was calculated and fed into further analyses as a marker of excitability-to-threat or dmPFC excitability regulation. To complement these measures, we computed the peak-to-peak differences, indicating the amplitude latency in between pairs of evoked components. For interpretational purposes we computed the TEPs using the same approach.

For our main study, the TMS experiment, we divided the scalp into frontal, dmPFC, occipital, central, parietal and temporal regions (supplementary Figure 1) and averaged data from the EEG channels from each of these regions.

### Heart rate estimation

The heart rate estimation was done from the EEG signals using the extended version of the independent component analysis (ICA) algorithm [25] based on information maximization [26].

For EEG analysis, the rows of the input matrix *y* are the EEG signals recorded at the 256 electrodes, the rows of the output data matrix *v* = *X y* are time courses of activation of the lCA components, and the columns of the inverse matrix, *X*^−1^, give the projection strengths of the respective components onto the scalp sensors.

In general, and unlike primary component analysis (PCA), the component time courses of activation will be non-orthogonal. Corrected EEG signals can then be derived as *y*′ = (*X*)^−1^*v*′ where *v*′ is the matrix of activation waveforms, *v*, with rows representing cardiac artefactual sources which are then extracted for further estimations from each participant. In total for the no-TMS experiment we concatenated the 36 CS+ trials without US to have 180 seconds and 24 CS-trials to have 120 seconds. The same was done for the TMS experiment.

### MRI data analysis

The individual MRI data was pre-processed using the FreeSurfer software package v5.3 (https://surfer.nmr.mgh.harvard.edu/). The automated pipeline [27, 28] included: (i) affine registration into Talairach space, (ii) intensity normalization for image inhomogeneities, (iii) removal of skull and non-brain tissues, (iv) definition of the gray/white matter and gray/cerebrospinal fluid boundaries, (v) surface creation and correction for topology defects and (vi) parcellation of the cortex and subcortical regions [29, 30].

Data of all subjects was visually inspected for errors during pre-processing and manually corrected when necessary. Volumes of the dmPFC and insula, amygdala and hippocampus of both cerebral hemispheres were computed and corrected by head size in a fully-automated fashion. The anatomical delimitation of the dmPFC was performed according to Etkin, Egner [31], see supplementary Figure 2.

**Figure 2.**
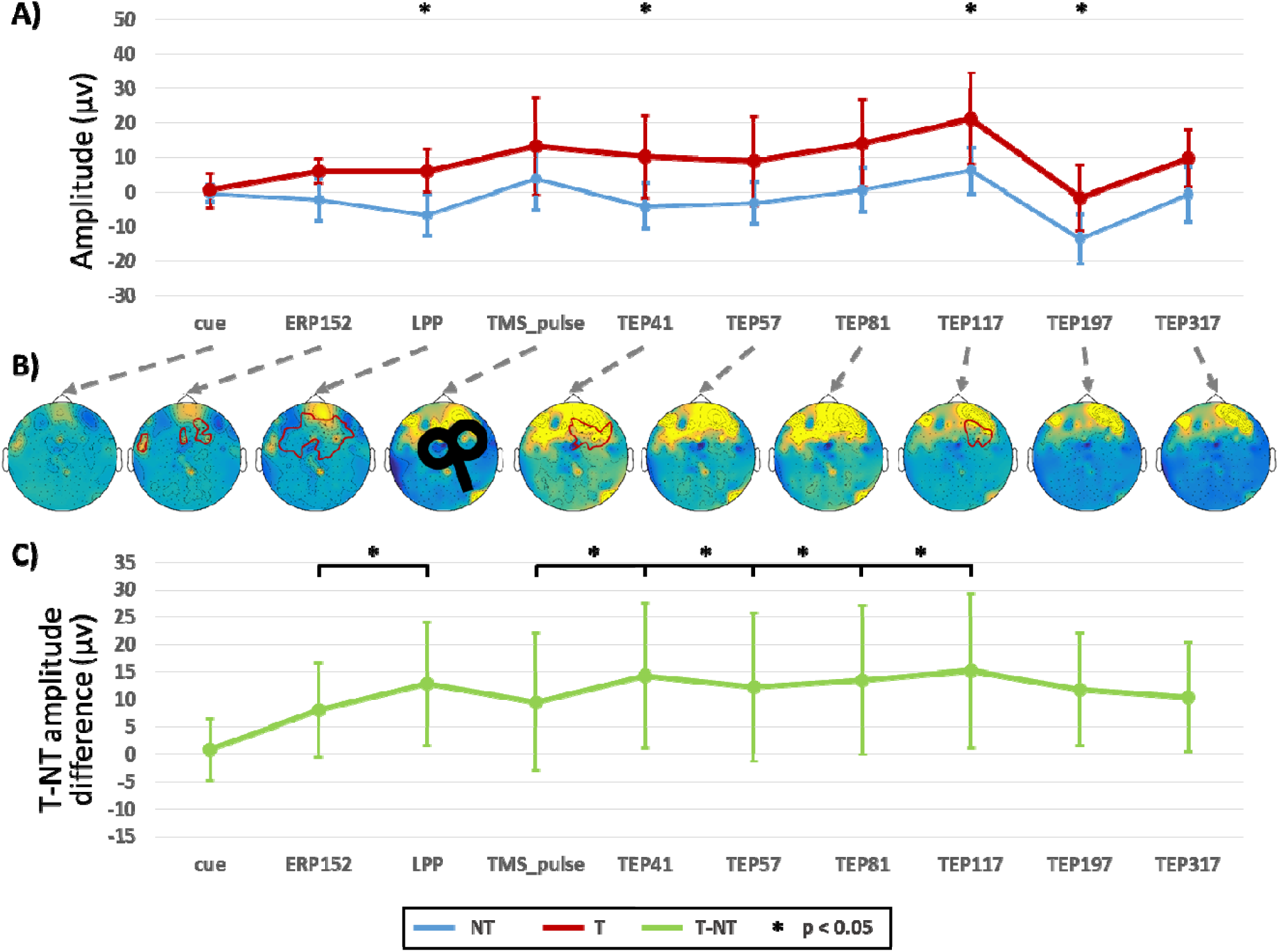
Event-related potentials (ERP) and TMS-evoked potentials (TEP). (A) Cortical excitability-to-threat (a.k.a. ERP) and dmPFC related modulation peak (a.k.a. TEP) amplitude differences between the threat (T) and no-threat (NT) conditions over scalp electrodes. (B) Topographical distribution of the amplitude differences across the scalp, the red lines show the significant electrodes after cluster analysis (Monte Carlo permutations, p < 0.05). (C) Peak-to-peak amplitude latency differences for the cortical excitability-to-threat and dmPFC related modulation in the main experiment. * denotes *p* < 0.05.

### Statistical analyses

The TMS-evoked responses over the dMPFC in the threat and no-threat conditions, as well as the fear ratings and registered heart rates, were compared using one-tailed paired t-tests. The effect sizes (d’), evaluated with the Cohen’s d, are reported for all comparisons. In order to evaluate the activity distribution of every evoked response while accounting for multiple comparisons, we applied non-parametric cluster-based statistics [32] with 1000 randomizations and a p-threshold of 0.05 to indicate channels within a cluster. After identification of significant amplitude differences, the dmPFC excitability-to-threat (see above) was extracted and further analyzed. Comparison of the cortical excitability-to-threat and dmPFC modulation in between the no-TMS and TMS data was carried out using t-test analyses.

Therefore, we adjusted multiple linear regression models to assess the predictive value of registered heart rates and the stress ratings to dmPFC excitability. The same models were used to investigate the anatomical substrates of the cortical excitability-to-threat and the dmPFC-related modulation. To avoid multicollinearity due to spurious correlations between different components of the evoked excitability-to-threat modulation, regression analyses were performed for each peak separately. Each excitability-to-threat peak was then regressed against volume of the hippocampus, amygdala, insula and dmPFC. We applied a backward elimination in the regression models, where predictors below a 10% significance level were deleted until none were left or statistical significance was reached.

Only supra-threshold values obtained after correction for multiple comparisons (FDR *p* < 0.05) for condition testing and after contrasting the regression slopes against the null hypothesis (F-test, *p* < 0.05) for the regression analyses were considered as significant and reported in the subsequent parts of the manuscript.

## Results

### Psycho-physiological markers of instructed fear responses

The subjective fear ratings, obtained from the questionnaires, showed that the T condition lead to higher levels of expectancy for threat (*p* < 0.001, d’ = 2.6) than NT cues. The captured heart rates were also markedly increased in T compared to the NT condition (*p* < 0.001, d’ = 1.98).

### Cortical excitability-to-threat evoked potentials and dmPFC excitability modulation

Concordant with the results on the subjective ratings and the heart rate frequency increase, the evoked peak amplitudes after the T visual cues were higher than for the NT trials in both the TMS and no-TMS experiments (Figure 2). More specifically, after the visual cues we could identify two amplitude peaks that showed modulation. The first ERP component occurred at 152 ms (ERP152), followed by the LPP with an increased and sustained activity with high amplitude at 500 ms. In the no-TMS experiment, the LPP showed a sustained response beyond 1 s, decaying at ∼1300 ms, whereas in the TMS experiment apart from the same two ERPs additional and differential modulation of the EEG activity after the TMS pulse was observed. Here, six TMS-evoked potentials have been identified, TEP1 at 41 ms, TEP2 at 57 ms, TEP3 at 81 ms, TEP4 at 117 ms, TEP5 at 197 ms and TEP6 at 317 ms. The topographical distribution of the excitability-to-threat and the corresponding peak amplitudes for both T and NT conditions in the TMS experiment are shown in Figure 2 (A, B).

When evaluating the evoked activity (Figure 2, Table 1), the amplitude of the ERP152 did not differ between the T and NT conditions. Whereas the LPP showed a significant increase in amplitude in the T condition (*p* = 0.038, Cohen’s d = 0.5). After the TMS pulse, three of the TEPs showed amplitude differences between the two conditions: TEP41 (*p* = 0.048, d’ = 0.47), TEP117 (*p* = 0.049, d’ = 0.47), TEP197 (*p* = 0.036, d’ =0.51). The cluster analyses showed significant increases in the T condition when compared to the NT condition in the sensors corresponding to the frontal, dmPFC and central regions (Figure 2 B).

**Table 1.** Differences between Threat and no-Threat conditions in the TMS experiment.

|  | t | p | Mean<br>Difference | SE<br>Difference | Cohen's<br>d |
| --- | --- | --- | --- | --- | --- |
| *Psycho-physiological markers of instructed fear responses* |  |  |  |  |  |
| Heart rates | 8.632 | **< .001** | 2.395 | 0.277 | 1.98 |
| Fear ratings | 11.73 | **< .001** | 0.465 | 0.04 | 2.623 |
| Cue (0 ms) | 0.269 | 0.395 | 0.78 | 2.898 | 0.06 |
| *Indicators of cortical excitability-to-threat and dmPFC modulation* |  |  |  |  |  |
| ERP152 | 1.86 | **0.039** | 8.096 | 4.353 | 0.416 |
| LPP | 2.235 | **0.019** | 12.891 | 5.769 | 0.5 |
| TMS pulse | 1.485 | 0.077 | 9.532 | 6.42 | 0.332 |
| TEP41 | 2.113 | **0.024** | 14.273 | 6.756 | 0.472 |
| TEP57 | 1.786 | **0.045** | 12.262 | 6.866 | 0.399 |
| TEP81 | 1.951 | **0.033** | 13.496 | 6.918 | 0.436 |
| TEP117 | 2.105 | **0.024** | 15.205 | 7.222 | 0.471 |
| TEP197 | 2.258 | **0.018** | 11.814 | 5.232 | 0.505 |
| TEP317 | 3.308 | **0.002** | 22.098 | 6.68 | 0.74 |
| *Peak-to-peak latencies* |  |  |  |  |  |
| 0-to-ERP152 | 1.153 | 0.132 | 7.316 | 6.343 | 0.258 |
| ERP152-to-LPP | 2.375 | **0.014** | 4.795 | 2.019 | 0.531 |
| LPP-to-TMS | 0.726 | 0.762 | 3.359 | 4.627 | 0.162 |
| TMS-to-TEP41 | 2.159 | **0.022** | 4.741 | 2.196 | 0.483 |
| TEP41-to-TEP57 | 1.829 | 0.958 | 2.011 | 1.099 | 0.409 |
| TEP57-to-TEP81 | 3.174 | **0.002** | 1.234 | 0.389 | 0.71 |
| TEP81-to-TEP117 | 2.361 | **0.015** | 1.708 | 0.724 | 0.528 |
| TEP117-to-TEP197 | 0.967 | 0.827 | 3.391 | 3.505 | 0.216 |
| TEP197-to-TEP317 | 0.866 | 0.801 | 1.387 | 1.602 | 0.194 |

t-tests contrast tested: Threat>no-Threat, bold text indicates significance ( $p < 0.05$ ).

**Table 2.**
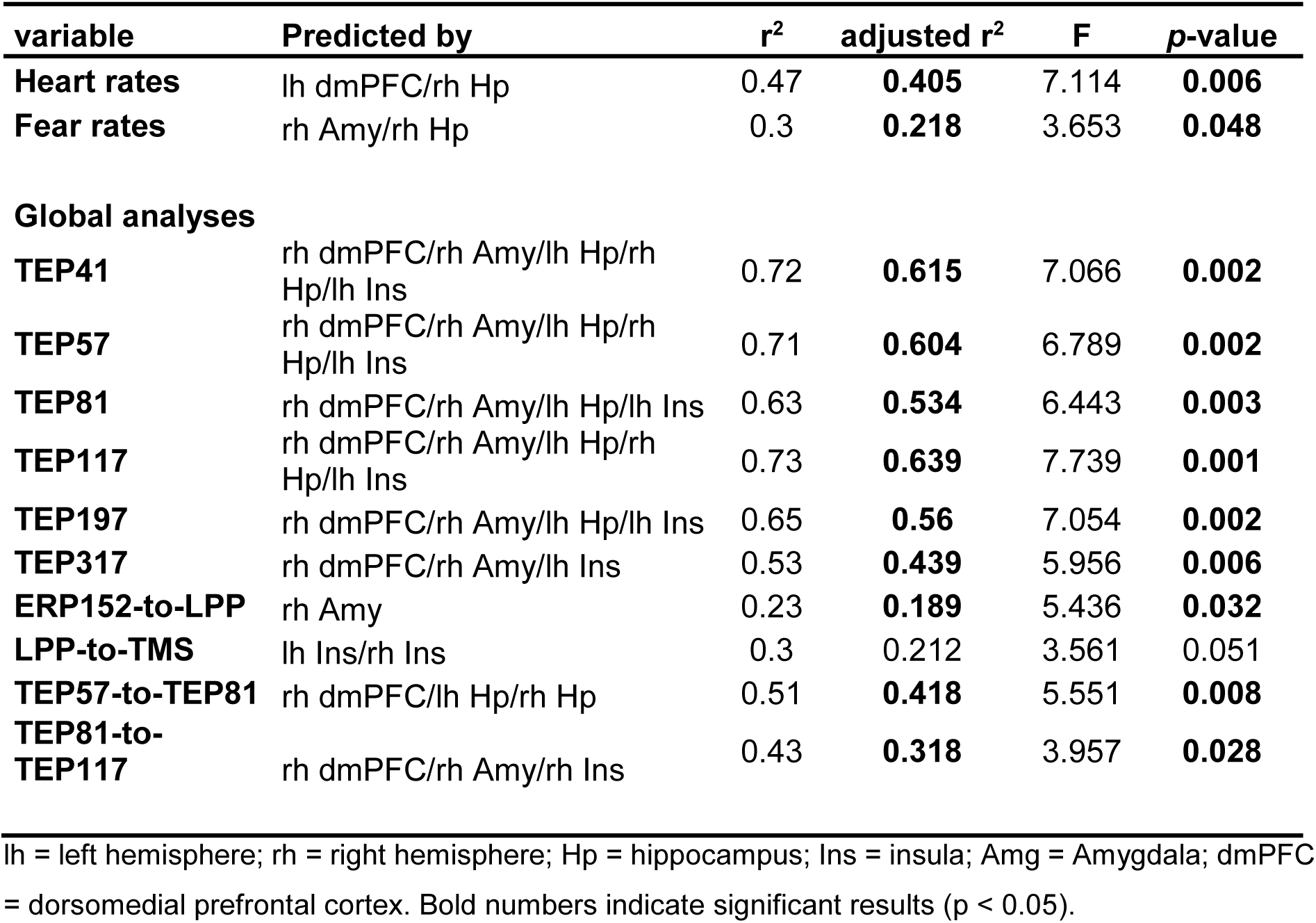
Backward step-regression analyses. lh = left hemisphere; rh = right hemisphere; Hp = hippocampus; Ins = insula; Amg = Amygdala; dmPFC= dorsomedial prefrontal cortex. Bold numbers indicate significant results (p < 0.05).

Henceforth, we used the cortical excitability-to-threat and the dmPFC-related modulation in the subsequent analyses (Figure 2 C). In the case of the peak-to-peak amplitude latencies, the LPP showed increased peak-to-peak amplitudes in respect to the ERP152 (*p* = 0.014, d’ = 0.73). After the TMS stimulation, four peak-to-peak latencies showed significant changes: increased TEP41 compared to the TMS onset (*p* = 0.022, d’ = 0.49), decreased TEP57 compared to TEP41 (*p* = 0.042; d’ = 0.41), increased TEP81 compared to TEP57 (*p* = 0.0025, d’ = 0.72) and increased TEP117 compared to TEP81 (*p* = 0.015, d’ = 0.58).

When comparing the markers of cortical excitability in the two experiments, no significant changes were detected for the cortical excitability to T. Of note, increased excitability was evidenced after the TMS stimulation over the dmPFC (Figure 3).

**Figure 3.**
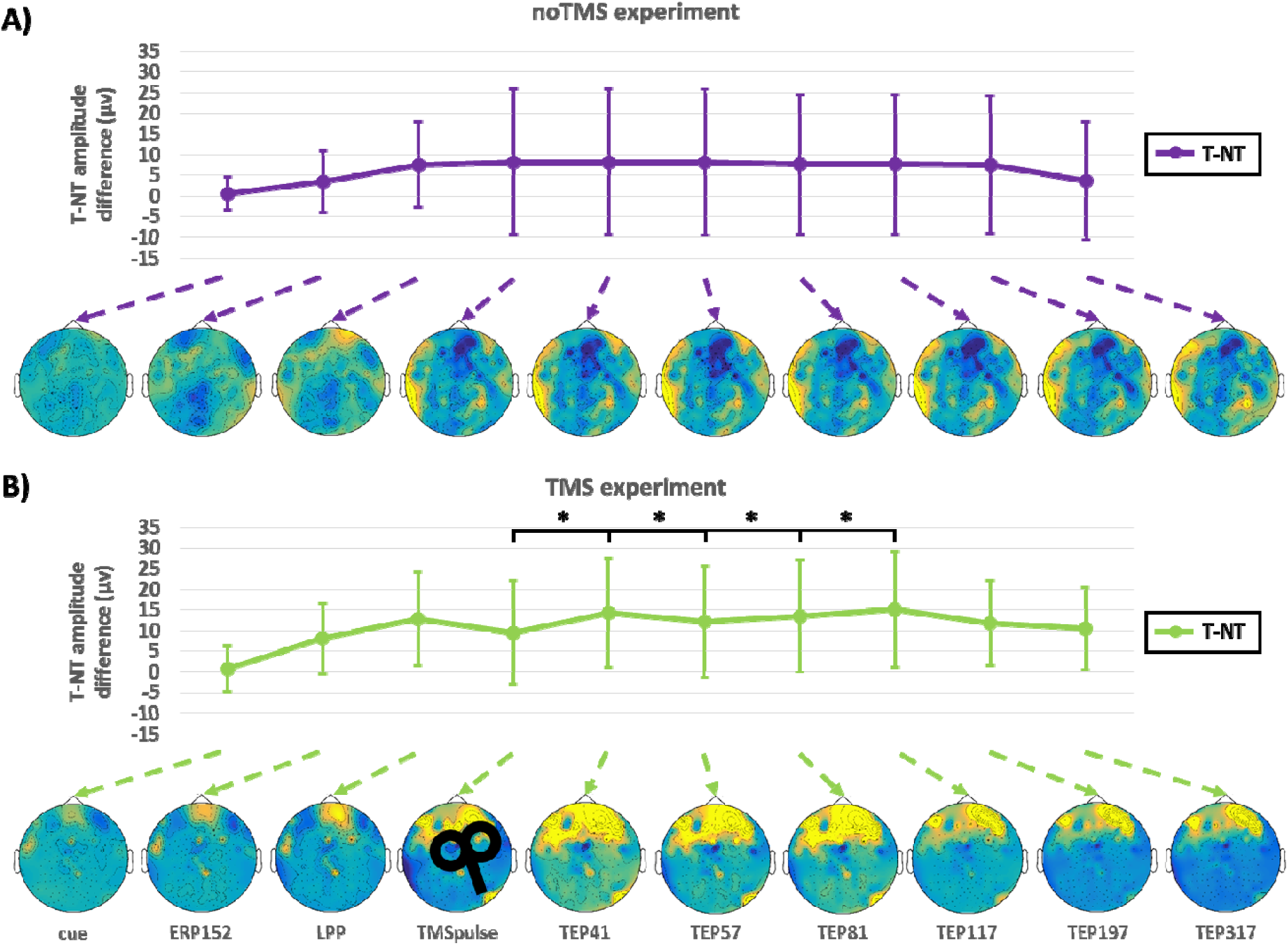
Comparison of cortical excitability-to-threat and the dmPFC related modulation between TMS and no-TMS experiments. T-NT differences and their topographical representations in the (A) no-TMS (purple) and (B) TMS (green) experiments. The black lines indicate significant peak-to-peak latency changes in the TMS experiment with respect to the no-TMS experiment. * denotes *p* < 0.05.

**Figure 4.**
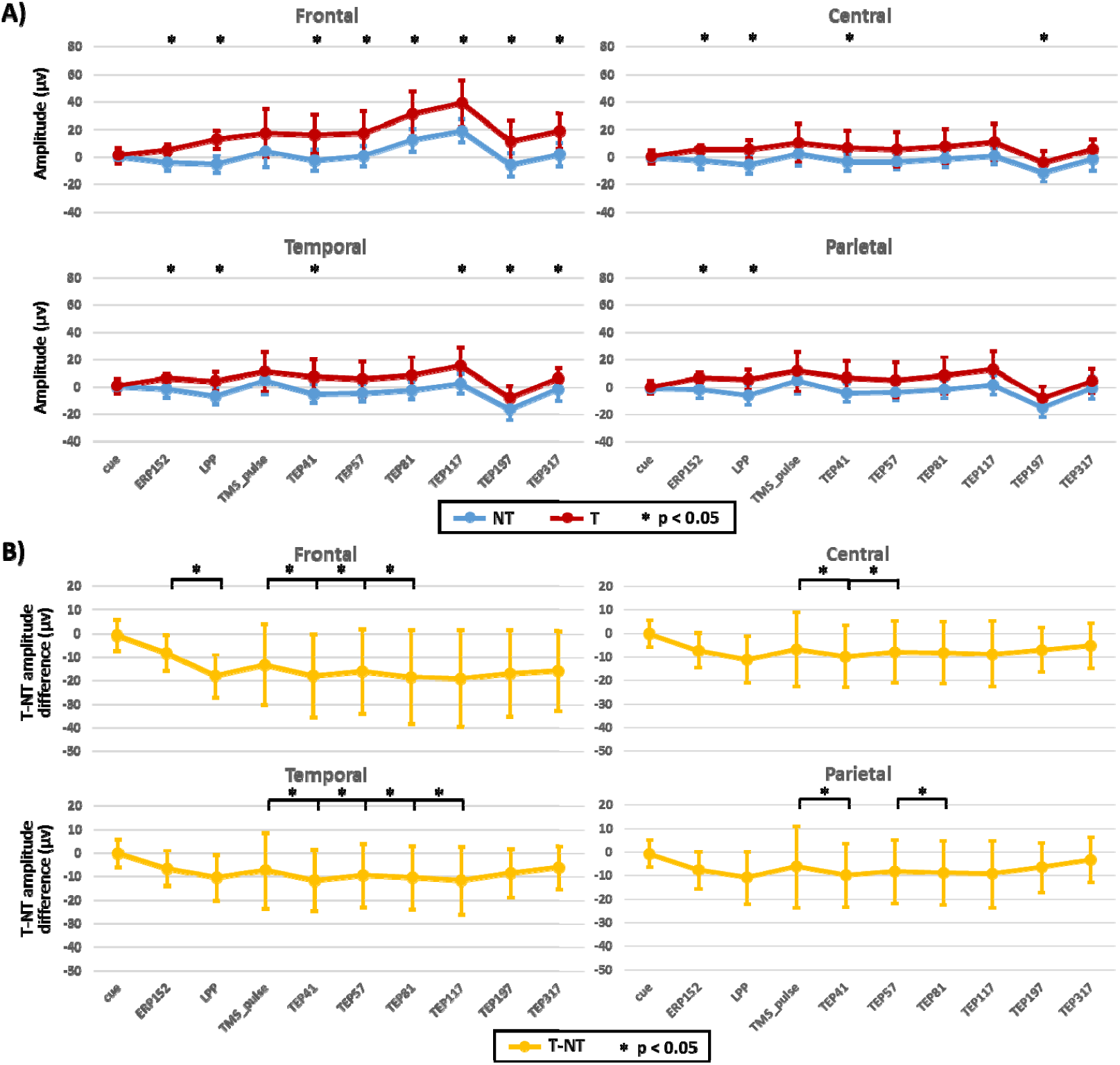
Regional event- and TMS-evoked potentials. (A) Amplitude peaks of the no-threat (NT) and threat (T) conditions. (B) Peak-to-peak latency activity of the excitability-to-threat and dmPFC-related modulation at different regions. * denotes *p* < 0.05.

### Structural substrates of the excitability-to-threat and dmPFC-related modulation

The fear ratings were predicted by both right hippocampus and amygdala volumes (*r*^*2*^= 0.3, F = 3.65; *p* = 0.048). Furthermore, the volume of the left dmPFC and the right hippocampus correlated with the increase in heart rates in response to fear (*r*^*2*^ = 0.47; F = 7.11; *p* = 0.006). As shown in Table 2, when examining associations between structural properties of the network nodes and the dmPFC excitability response to T, the integrity of the right dmPFC and the right amygdala predicted these measures. When analyzing the excitability-to-threat at the regional level, the structural integrity of the right dmPFC predicted the amplitudes of the evoked potentials after TMS. A similar effect was shown as well for several other structures in the network (i.e. insula, see also supplementary Table 1 for more detailed information).

## Discussion

In this work, we characterize excitability regulation patterns during an instructed fear paradigm non-invasively in healthy subjects. Threat processing is related to increased event-related activity with topological maximum in the dMPFC area. Furthermore, TMS pulses over the dMPFC induced a consistent modulation of the event-related activity with longer and increased threat-related responses. The threat-dependent excitability modulation was linked to the LPP component of evoked response. We found highly significant associations between evoked responses and markers of gray matter integrity, mainly in the dmPFC, but also in the amygdala and insula. The applied integrative approach delineates illustratively the involved network. Our findings add to the current literature showing a pivotal role of the dmPFC in controlling the adaptive fear responses and introduce a non-invasive paradigm to measure physiological responses to threat [2]. Furthermore, the strong interrelation of excitability fingerprints, microstructural integrity and physiological markers of fear processing such as heart rate underpins the pivotal integrative value of the dmPFC in evaluative processes related to threat and the robust value of the introduced paradigm for causal interrogation of this specific node for fear processing.

The applied EEG approach permits an exact temporal characterization of threat processing. We see no delimitation of fear processing at early phases as quantified by ERP responsiveness at 152 ms (ERP152). And indeed at this very early stage merely the processing of complex visual information occurs and not the difference in the valence of the stimuli [33]. In our study, in both the TMS and no-TMS experiments increased threat-related excitability, as reflected by the appearance (and increase) of the LPP component, was detected. LPP is characterized by a sustained activity from ∼500 ms and beyond ∼1000 ms. Stimuli, most directly relevant to biological or imperative contents (threat, mutilation, etc), lead to an increase of LPP [34, 35]. Emotional modulation of the LPP lasts for the full duration of stimulus presentation and thereafter (e.g., 300–1500 ms) and show several topographic shifts from parietal to central and frontal representations [4, 36, 37]. Moreover, LPP seems not to habituate to emotional stimuli. Amygdala and prefrontal cortex activation have been described and related to attentional processing of emotional stimuli [38]. However, LPP amplitudes have been clearly shown to be linked to memory encoding and storage [39-43]. Similar to existing data showing poor correlation of the magnitude of early components (ERP152) with threat encoding, we only see consistent differences in the T-NT processing in the late components indexing distinct temporal facets of threat processing [9, 39]. According to these finding, the LPP likely represents the summated activity of the entire network, thus through specific causal manipulation, possible by optogenetics in animal studies or TMS in humans, a specific role of each node of the fear network could be delimitated.

In our study, the evoked responses are further modulated by TMS pulses over the dMPFC. This area might mediate the explicit evaluation of fear states and grant a controlled processing [16, 44]. In concordance, larger LPP responses and increased dmPFC activations have been associated with amplified states of fear [2, 45]. LPP increase and prolongation might mirror increased processing of the threat stimuli, while the dmPFC interferes with these processes.

TMS pulses induce well-described evoked potentials [46, 47], however little is known about TMS modulation of the ongoing activity over cortical areas other than the motor cortex [48]. TEPs reflect the excitability of the cortex at the stimulated area and represent a summation of excitatory and inhibitory phenomena [49]. A recent work addressed TEP over prefrontal areas showing mainly an inhibitory effect of single-pulse TMS over prefrontal cortex [46]. However, the evoked activity is more a complex interplay with possible interactions with the excitatory neurotransmission for the early TEP peaks (10–30 ms) and inhibitory phenomena at later peaks (100–200 ms) as known for motor cortex stimulation [48]. Moreover, every TEP component presents a different topology suggesting a dynamic interaction of distinct cortical and subcortical areas.

In our study, the peak latency from ERP152-to-LPP is predicted by the volume of the right amygdala. Furthermore, the subjective fear rates were predicted by the integrity of the right amygdala. Both findings support the role of amygdala for fear evaluation and processing. These results are in good agreement with pivotal studies, showing that the activity of the amygdala was positively correlated with subjective reports of anxiety [50]. The amygdala innervates the autonomic system, and thus is involved in the modulation of physiological responses to threat, aversive stimuli and signs of anxiety arousal, such as changes in heart rate [2, 51]. Furthermore, the structural integrity of the paths connecting the amygdala to frontal regions predicts anxiety levels [15]. Similar to our study, recent work showed that threat events enhance dmPFC-amygdala connectivity [52], while the dmPFC possibly modulates amygdala activation, probably guided by inhibitory projections [53, 54]. Here, these mechanisms were evidenced by both structural MRI and ERP analyses, highlighting not only a critical role of the dmPFC in regulating threat-related excitability, but also in recruiting interconnected regions of the fear network.

### Clinical and functional relevance

Contrary to Pavlovian conditioning, in the addressed fear paradigms the subjects are instructed before the experiment about T and NT conditions. Hence, for fear conditioning, learning takes place prior to stimuli exposure and fear processing requires controlled evaluation of fearful stimuli [55], likely with involvement of different areas of the fear network. Previous studies on instructed fear have consistently shown activations of the dmPFC, amygdala and ACC [2], while robust measures that are easy to apply in the clinical setting are still lacking. The dmPFC is not directly involved in the initial generation of fear responses but specifically modulate controlled/attended threat processing. Indeed, it has been shown that a loss of function in the dorsal mPFC regions is related with prolonged amygdala activation in persons with emotional dysregulation [14]. Prefrontal cortex excitability abnormalities have been linked to impaired threat processing, anxiety and depression as well [56-58]. Taken together, these findings suggest that measures of dmPFC responses could evolve into a translational fingerprint that could be applied in experimental or clinical settings to dissect physiological from pathological responses or monitor the transitional dynamics which provide resilience to mental illness. Furthermore, our results highlight the link between anatomical integrity and brain excitability patterns.

## Conclusions

According to previous studies showing increased activation of the dmPFC during fear processing, the current paradigm allowed us to examine the specific role of dmPFC excitability regulation in healthy young subjects. TMS pulses over right dmPFC during the instructed fear paradigm induce evoked responses with distinct temporal patterns linked to structural nodes properties of the fear network. Providing conclusive evidence for the involvement of the dmPFC in modulating the excitability-to-threat related to the long-lasting LPP component, a marker of fear stimuli processing, accompanied by predictions of structural integrity for different TMS-evoked activity peaks. Our results show causal evidence that fear processing requires higher cognitive mechanisms guided by the excitability properties of the dmPFC. These paradigm can be applied to test specific effects of dmPFC excitability modulation related to resilience and health.

## Acknowledgments

The authors thank Cheryl Ernest for proofreading the manuscript. The authors gratefully acknowledge the computing time granted on the supercomputer Mogon at Johannes Gutenberg University Mainz (hpc.uni-mainz.de).

Funding: This work was supported by a grant from the German Research Council (DFG; CRC-TR-1193).

