## Supplementary Materials for "Excitability regulation in the dorsomedial prefrontal cortex during sustained instructed fear responses: a TMS-EEG study"

### Supplementary material

**Supplementary Figure 1.** Regional aggrupation of the EEG sensors according to the underlying brain regions (lobules).

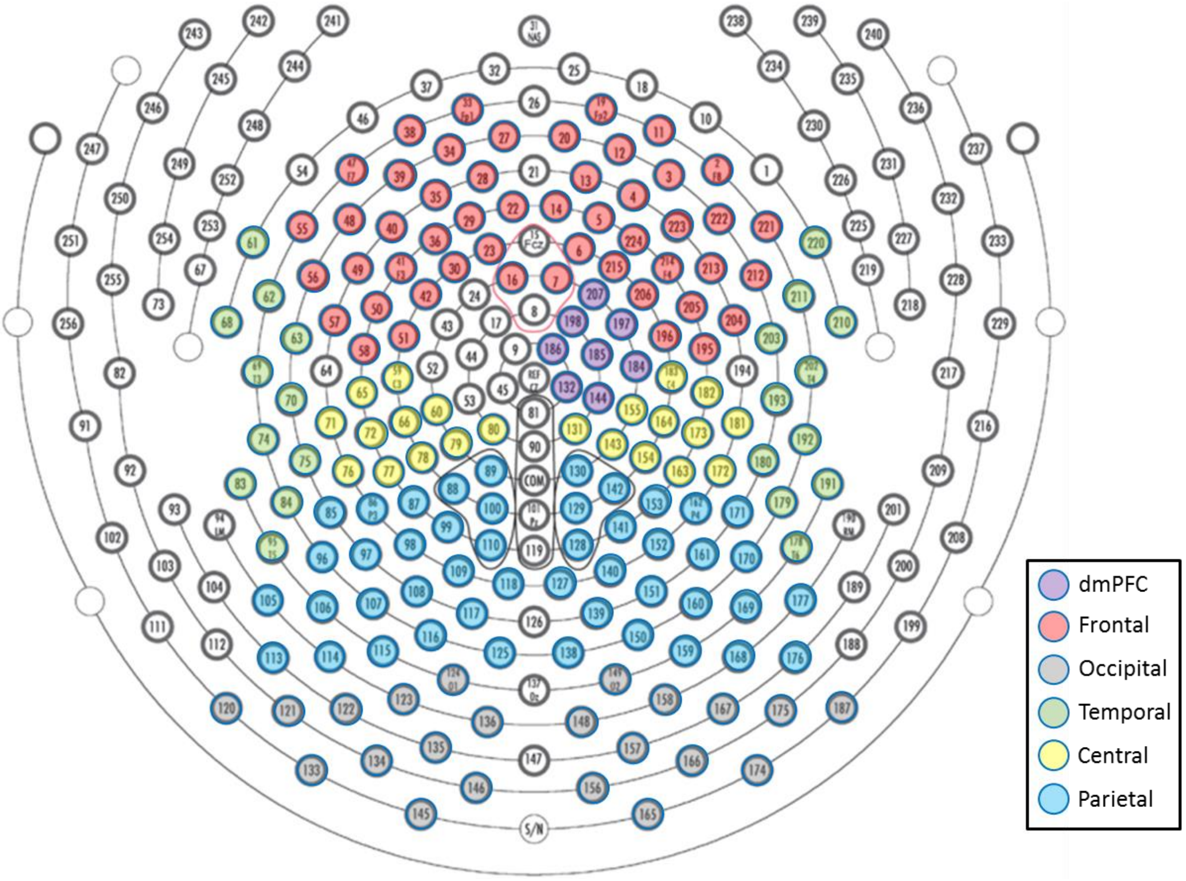

**Supplementary Figure 2.** Parcellation of the medial prefrontal cortex (mPFC) according to Etkin et al., (2011). Abbreviations: ACC, anterior cingulate cortex; sg, subgenual; pg, pregenual; vm, ventromedial; rm, rostromedial; dm, dorsomedial; ad, anterior dorsal; pd, posterior dorsal.

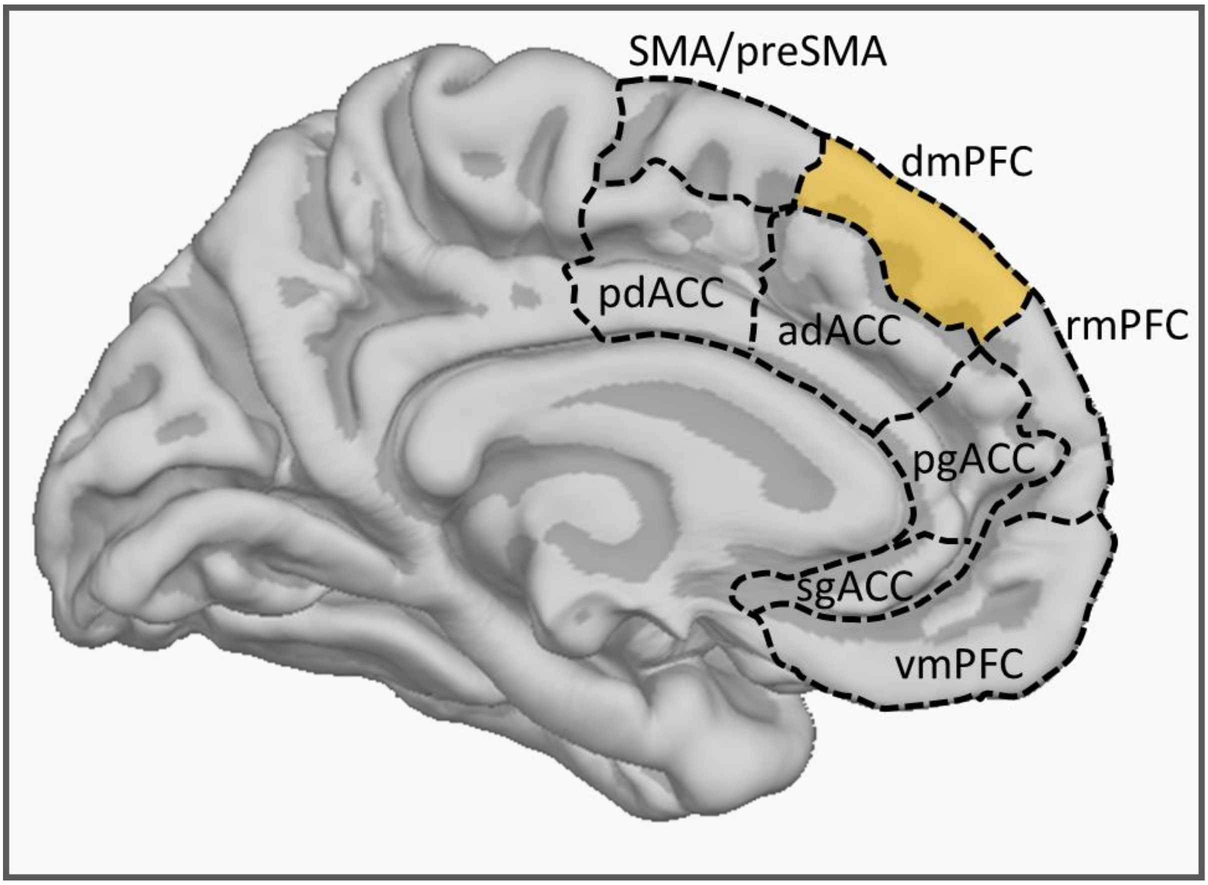

**Supplementary Figure 3.** A) Hear rates and B) subjective fear ratings of the no-TMS and TMS experiments. The red bar represents T and blue bar represents NT conditions. * denotes $p < 0.05$.

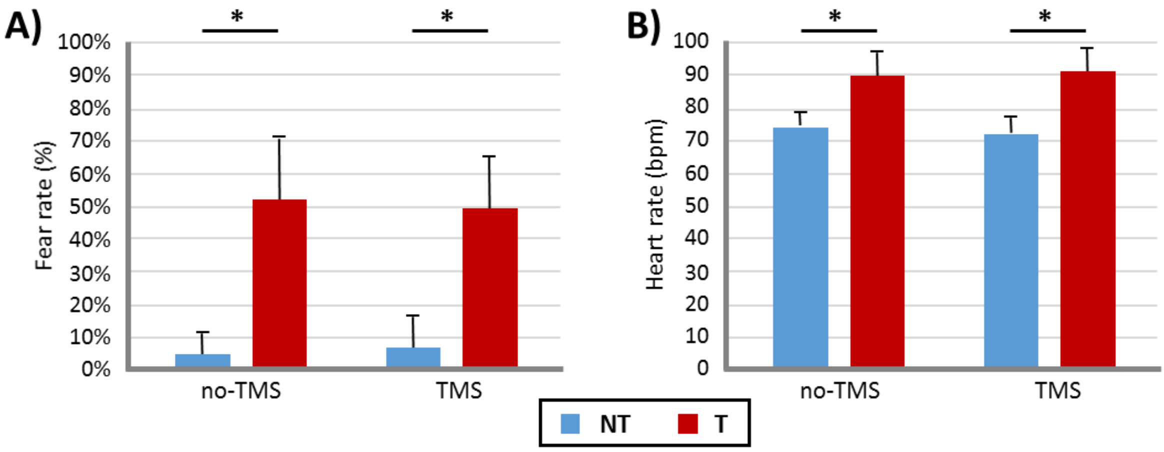

**Supplementary Table 1.** Backward step-regression analyses on the sensors grouped by the underlying brain regions. Significant predictions are shown ($p$ < 0.05).

| variable | Predicted by | $r^2$ | adjusted $r^2$ | F | *p*-value |
|---|---|---|---|---|---|
| **Frontal-LPP** | lh Hp/rh Hp/rh Ins | 0.52 | 0.434 | 5.852 | **0.007** |
| **dmPFC-TEP41** | lh Ins/lh Hp | 0.34 | 0.263 | 4.392 | **0.029** |
| **dmPFC-TEP57** | lh Ins/lh Hp | 0.4 | 0.334 | 5.757 | **0.012** |
| **dmPFC-TEP81** | lh Ins/lh Hp | 0.39 | 0.322 | 5.517 | **0.014** |
| **dmPFC-TEP117** | rh dmPFC/rh Amy/lh Hp/lh Ins | 0.59 | 0.48 | 5.393 | **0.007** |
| **dmPFC-TEP197** | rh dmPFC/rh Amy/lh Hp/lh Ins | 0.77 | 0.711 | 12.66 | < **.001** |
| **dmPFC-TEP317** | rh dmPFC/rh Amy | 0.5 | 0.444 | 8.571 | **0.003** |
| **Frontal-TEP41** | rh dmPFC/rh Amy/lh Ins/lh Hp/rh Hp | 0.64 | 0.505 | 4.882 | **0.009** |
| **Frontal-TEP57** | rh dmPFC/rh Amy/lh Ins/lh Hp/rh Hp | 0.67 | 0.549 | 5.62 | **0.005** |
| **Frontal-TEP81** | rh dmPFC/rh Amy/lh Ins/lh Hp/rh Hp | 0.64 | 0.515 | 5.033 | **0.008** |
| **Frontal-TEP117** | rh dmPFC/rh Amy/lh Ins/lh Hp/rh Hp | 0.66 | 0.536 | 5.387 | **0.006** |
| **Frontal-TEP197** | rh dmPFC/lh Ins/lh Hp | 0.43 | 0.323 | 4.027 | **0.026** |
| **Occipital-TEP41** | rh dmPFC/lh Ins/lh Hp | 0.49 | 0.399 | 5.199 | **0.011** |
| **Occipital-TEP57** | rh dmPFC/lh Ins/lh Hp | 0.43 | 0.322 | 4.002 | **0.027** |
| **Occipital-TEP81** | rh dmPFC/lh Ins/lh Hp | 0.43 | 0.327 | 4.072 | **0.025** |
| **Occipital-TEP117** | rh dmPFC/lh Ins/lh Hp | 0.47 | 0.375 | 4.803 | **0.014** |
| **Occipital-TEP197** | rh dmPFC/rh Amy | 0.48 | 0.413 | 7.685 | **0.004** |
| **Occipital-TEP317** | rh dmPFC/rh Amy/lh Hp | 0.65 | 0.589 | 10.08 | < **.001** |
| **Central-TEP41** | rh dmPFC/rh Amy/lh Hp/lh Ins | 0.59 | 0.486 | 5.484 | **0.006** |
| **Central-TEP57** | rh dmPFC/rh Amy/lh Ins/rh Ins | 0.58 | 0.468 | 5.182 | **0.008** |
| **Central-TEP81** | rh dmPFC/rh Amy/lh Ins/rh Ins | 0.59 | 0.477 | 5.325 | **0.007** |
| **Central-TEP117** | rh dmPFC/rh Amy/lh Ins/rh Ins | 0.62 | 0.514 | 6.018 | **0.004** |
| **Central-TEP197** | rh dmPFC/rh Amy/lh Hp/lh Ins | 0.77 | 0.708 | 12.51 | < **.001** |
| **Central-TEP317** | rh dmPFC/rh Amy/lh Hp/lh Ins | 0.74 | 0.666 | 10.46 | < **.001** |
| **Parietal-TEP41** | lh Ins/lh Hp | 0.38 | 0.301 | 4.735 | **0.015** |
| **Parietal-TEP57** | rh dmPFC/rh Amy/lh Ins/rh Ins | 0.51 | 0.377 | 3.879 | **0.023** |
| **Parietal-TEP81** | rh dmPFC/rh Amy/lh Ins/rh Ins | 0.51 | 0.375 | 3.855 | **0.024** |
| **Parietal-TEP117** | rh dmPFC/rh Amy/lh Hp/lh Ins | 0.58 | 0.467 | 5.167 | **0.008** |
| **Parietal-TEP197** | rh dmPFC/rh Amy | 0.54 | 0.484 | 9.921 | **0.001** |
| **Parietal-TEP317** | rh dmPFC/rh Amy/rh Hp | 0.71 | 0.656 | 13.08 | < **.001** |
| **Temporal-TEP41** | rh dmPFC/rh Amy/lh Hp/lh Ins | 0.66 | 0.57 | 7.299 | **0.002** |
| **Temporal-TEP57** | lh Ins/lh Hp | 0.45 | 0.385 | 6.936 | **0.006** |
| **Temporal-TEP81** | rh dmPFC/rh Amy/lh Hp/lh Ins | 0.61 | 0.509 | 5.916 | **0.005** |
| **Temporal-TEP117** | rh Amy | 0.67 | 0.581 | 7.599 | **0.001** |
| **Temporal-TEP197** | rh dmPFC/rh Amy/lh Hp/lh Ins | 0.74 | 0.665 | 10.44 | < **.001** |
| **Temporal-TEP317** | rh dmPFC/rh Amy/lh Hp/lh Ins | 0.7 | 0.626 | 8.939 | < **.001** |

lh = left hemisphere; rh = right hemisphere; Hp = hippocampus; Ins = insula; Amg = Amygdala; dmPFC = dorsomedial prefonral cortex. Bold numbers indicate significant results (p < 0.05).
